# Translational buffering by ribosome stalling in upstream open reading frames

**DOI:** 10.1101/2022.01.06.475296

**Authors:** Ty Bottorff, Adam P. Geballe, Arvind Rasi Subramaniam

**Affiliations:** Basic Sciences Division and Computational Biology Section of the Public Health Sciences Division, Fred Hutchinson Cancer Research Center, Seattle, WA 98109, USA; Biological Physics, Structure and Design Graduate Program, University of Washington, Seattle, WA 98195, USA; Human Biology and Clinical Research Divisions, Fred Hutchinson Cancer Research Center, Seattle, WA 98109, USA

## Abstract

Upstream open reading frames (uORFs) are present in over half of all human mRNAs. uORFs can potently regulate the translation of downstream open reading frames by several mechanisms: siphoning away scanning ribosomes, regulating re-initiation, and allowing interactions between scanning and elongating ribosomes. However, the consequences of these different mechanisms for the regulation of protein expression remain incompletely understood. Here, we performed systematic measurements on the uORF-containing 5’ UTR of the cytomegaloviral *UL4* mRNA to test alternative models of uORF-mediated regulation in human cells. We find that a terminal diproline-dependent elongating ribosome stall in the *UL4* uORF prevents decreases in main ORF translation when ribosome loading onto the mRNA is reduced. This uORF-mediated buffering is insensitive to the location of the ribosome stall along the uORF. Computational kinetic modeling based on our measurements suggests that scanning ribosomes dissociate rather than queue when they collide with stalled elongating ribosomes within the *UL4* uORF. We identify several human uORFs that repress main ORF translation via a similar terminal diproline motif. We propose that ribosome stalls in uORFs provide a general mechanism for buffering against reductions in main ORF translation during stress and developmental transitions.

## Introduction

About half of human mRNAs have at least one upstream open reading frame (uORF) in their 5’ untranslated region^1–3^. Ribosome profiling studies estimate that at least twenty percent of these uORFs are actively translated^4,5^. uORFs can regulate gene expression via the biological activity of the uORF peptide, but they also often *cis*-regulate translation of the downstream main ORF^6,7^. Despite having poor initiation sequence contexts, many eukaryotic uORFs repress main ORF translation^1,3,4,7-11^. uORF mutations are implicated in several human diseases via changes to main ORF translation^12,13^. For example, uORF mutations in oncogenes and tumor suppressors can act as driver mutations in cancer^14,15^.

uORFs can regulate translation via a variety of molecular mechanisms. uORFs can constitutively repress translation by siphoning away scanning ribosomes from initiating at downstream main ORFs. Multiple uORFs can interact together to regulate the re-initiation frequency at the main ORF. For example, uORFs in the *GCN4* (*S. cerevisiae* homolog of human *ATF4*) mRNA render main ORF translation sensitive to cellular levels of the eIF2α-GTP-tRNA_Met_ ternary complex^16,17^. Although initiation rate usually limits translation,^18–20^ inefficient elongation or termination on uORFs can regulate translation by barricading scanning ribosomes from reaching the main ORF^21–26^. Inefficient elongation can be driven by the nascent uORF peptide^27,28^, poorly translated codons in the uORF^29,30^, or small molecules such as amino acids or polyamines^23,24^. Further, interactions between scanning and elongating ribosomes on uORFs may cause dissociation of scanning ribosomes or enhanced initiation at start codons^23,31,32^.

Despite the plethora of proposed uORF regulatory mechanisms, their implications for the regulation of protein expression are not clear. For example, are some uORF regulatory mechanisms more effective than others at repressing protein expression across a wide range of biochemical parameters? How do uORFs alter the response of main ORF translation to changes in cellular and environmental conditions? Answering these questions requires a joint accounting of how the different steps of translation, such as initiation, scanning, and elongation, together influence the overall rates of uORF and main ORF translation. Since it is not straightforward to monitor the rates of individual steps of translation^33^, indirect measurements of protein expression are often necessary to infer the underlying mechanism of uORF-mediated regulation. Such inference requires rigorous kinetic models of uORF regulation that make testable experimental predictions for the effects of genetic mutations on protein expression.

Computational kinetic modeling has been widely used to study mechanisms of translational control^34^. Quantitative modeling of uORF translation has been used to support the regulation re-initiation model for the *GCN4* mRNA^35–37^. A computational model predicted that elongating ribosomes can dislodge leading scanning ribosomes on uORFs and confer stress resistance to protein expression^38^. However, these models have not been compared against alternative models of uORF regulation that predict queuing or dissociation of scanning ribosomes that collide with paused elongating ribosomes^21,23^. A critical barrier for such comparison has been the lack of a computational framework for the specification and simulation of different kinetic models of uORF-mediated translational regulation. Such a computational framework is necessary for the identification of unique experimental signatures of each proposed model and their comparison with experimental measurements. Even though simulation code has been made available in many computational studies of mRNA translation^18,38^, it is often highly tailored for specific models and cannot be easily modified to consider alternative regulatory mechanisms.

Here, we use experimental measurements on the well studied uORF-containing 5’ UTR of the human cytomegalovirus *UL4* gene to test different kinetic models of uORF-mediated translational control^21^. The second uORF (uORF2 henceforth) in the *UL4* 5’ UTR contains a terminal diproline motif that stalls 80S ribosomes by disrupting peptidyl transferase center activity^27,28^. For systematic model comparisons, we rely on a recent computational framework that allows easy specification and efficient simulation of arbitrary kinetic models of translational control^39^. Using this experimentally-integrated modeling approach, we find that the presence of 80S stalls in uORF2 of *UL4* 5’ UTR confers resistance (called buffering henceforth) of main ORF translation to reduced ribosome loading on the mRNA. Modeling suggests that collisions of scanning ribosomes with the stalled 80S ribosome confer this buffering behavior. Experimental variation of the distance between the uORF2 start codon and elongating ribosome stall supports a kinetic model in which scanning ribosomes dissociate rather than queue upon colliding with the 80S stall. We also identify several human uORFs that have repressive terminal diproline motifs similar to the *UL4* uORF2 80S stall. We propose that ribosome stalls in uORFs enable buffering of main ORF translation against reduced ribosome loading across cellular and environmental transitions. Together, our results illustrate the value of experimentally-integrated kinetic modeling for comparisons of different uORF regulatory mechanisms and identification of novel experimental signatures from complex molecular interactions.

## Results

### Models of uORF regulation of main ORF translation

We surveyed five previously proposed models of uORF regulation of main ORF translation (Fig. 1). We distinguished between these models using a combination of computational modeling and experimental reporter assays. In the constitutive repression model9 (Fig. 1A), uORFs siphon away scanning ribosomes from the main ORF since re-initiation is usually infrequent^40–43^. In the 80S-hit dissociation model^38^ (Fig. 1B), elongating ribosomes that hit downstream scanning ribosomes cause the 3’ scanning ribosomes to dissociate from the mRNA. In the queuing-mediated enhanced repression model^23^ (Fig. 1C), a stalled elongating ribosome within the uORF allows upstream scanning ribosomes to queue in the 5’ region. This queuing can bias scanning ribosomes to initiate translation at the uORF rather than leaky scan past it. In the collision-mediated 40S dissociation model^31,32^ (Fig. 1D), scanning ribosomes instead dissociate if they collide with a 3’ stalled elongating ribosome.

**Figure 1.**
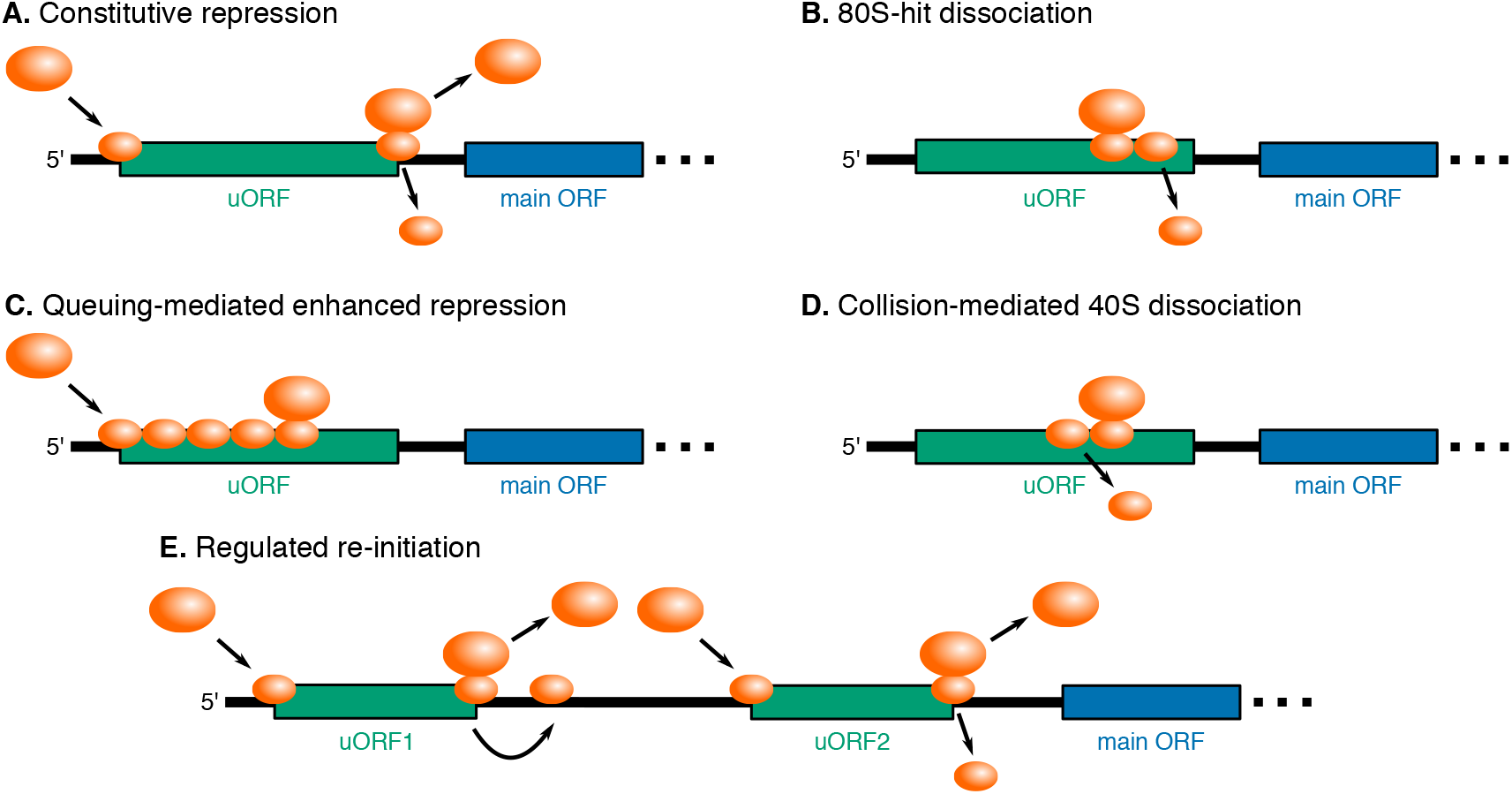
Models of uORF regulation considered in this study. **(A)** *Constitutive repression.* The uORF constitutively siphons away a proportion of scanning ribosomes from the main ORF. **(B)** *80S-hit dissociation.* Elongating ribosomes that collide with 3’ scanning ribosomes cause the leading scanning ribosome to dissociate from the mRNA. **(C)** *Queuing-mediated enhanced repression.* Scanning or elongating ribosomes form a queue behind a 3’ stalled elongating ribosome. If the queue correctly positions a scanning ribosome at the uORF start codon, then the proportion of scanning ribosomes that initiate translation at the uORF increases. **(D)** *Collision-mediated 40S dissociation.* Scanning ribosomes that collide with a 3’ stalled elongating ribosome dissociate from the mRNA. **(E)** *Regulated re-initiation.* Ribosomes initiate translation at the first uORF, and scanning continues after termination. Ribosomes reinitiate at the main ORF or the second downstream uORF when phosphorylated eIF2α levels are high or low, respectively. The schematic is depicted in a low phosphorylated eIF2α state.

Lastly, in the regulated re-initiation model^16,44,45^ (Fig. 1E), for example in the *GCN4* (*S. cerevisiae* homolog of human *ATF4*) mRNA, translation of the first uORF is followed by re-initiation at either a second downstream uORF or the main ORF depending on the stress status of the cell. After terminating at the first uORF, scanning ribosomes must reacquire a new eIF2α-GTP-tRNA_Met_ ternary complex (TC) before re-initiating. The time to reacquire a new TC correlates with the proportion of phosphorylated eIF2α. Therefore, when cells are not stressed and the proportion of phosphorylated eIF2α is lower, translation of the first uORF is followed by reinitiation at the second downstream uORF. Alternatively, when cells are stressed and the proportion of phosphorylated eIF2α is higher, translation of the first uORF is instead followed by re-initiation at the main ORF.

### Experimental system for testing different models of uORF-mediated translational regulation

To differentiate between proposed models of uORF regulation (Fig. 1), we used the well studied human cytomegaloviral *UL4* uORF231 as an experimental model. The 5’ leader region preceding the *UL4* coding sequence contains three uORFs (Fig. 2A). uORF1 slightly reduces uORF2 repressiveness by siphoning scanning ribosomes away from uORF2, and uORF3 is irrelevant for repression^31^. uORF2 represses main ORF translation via a terminating ribosome stall that is dependent on the uORF2 peptide sequence21 (Fig. 2A, irrelevant uORFs boxed in white, key uORF2 boxed in green). The two C-terminal proline residues in uORF2 are necessary for the elongating ribosome stall. These residues are poor substrates for nucleophilic attack to generate a peptide bond and also reorient the ribosomal peptidyl transferase center to reduce termination activity^28^. Reduced termination activity is further mediated through an interaction between the uORF2 nascent peptide and the GGQ motif within eRF1^46^. Even though the A-site of the uORF2-stalled ribosome is occupied by a stop codon, we will refer to it as an elongating ribosome stall since they are functionally equivalent for the purposes of this study. This terminology is also inclusive of elongation stalls within other uORFs^22,23,47–51^.

**Figure 2.**
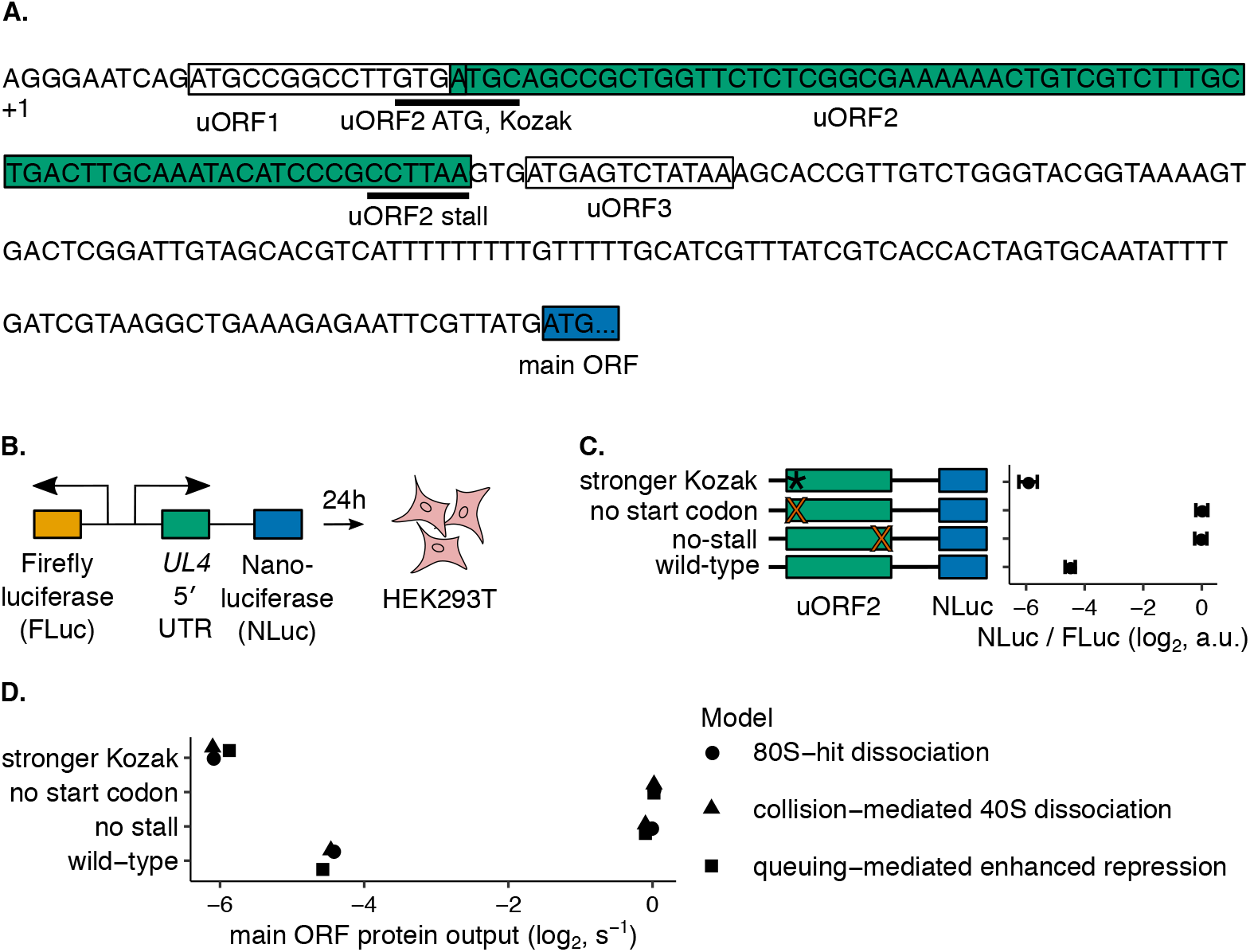
An experimental and computational platform for assessing uORF-mediated regulation of main ORF translation. **(A)** The 236 nt 5’ UTR of *UL4* mRNA from human cytomegalovirus contains 3 uORFs. The terminal proline and stop codons of uORF2 at which the P- and A-sites of the stalled ribosome are positioned is highlighted as uORF2 stall. **(B)** A dual-luciferase reporter system measures 5’ UTR repressiveness in HEK293T cells. Firefly luciferase signal serves as an internal control for transient transfection efficiency. **(C)** The reporter system recapitulates the known elongating ribosome stall-dependent repression of translation by the *UL4* uORF2^21^. The indicated mutations improve the uORF2 Kozak context (ACCATGG instead of GTGATGC), remove the start codon (ACC instead of ATG), or remove the elongating ribosome stall by mutating the terminal proline to alanine codon (GCT instead of CCT). Error bars show standard error of mean NLuc / FLuc ratios over 3 technical replicates. Data are normalized to a no-uORF start codon control. **(D)** Computationally predicted uORF regulation in the 80S-hit dissociation, queuing-mediated enhanced repression, and collision-mediated 40S dissociation models. Data are normalized to a no-uORF start codon control. The parameter combination that best recapitulated the control behavior in Fig. 2C is displayed in Table 1. Error bars of simulated data are smaller than data points.

We inserted the uORF2-containing *UL4* leader sequence into a dual-luciferase reporter system (Fig. 2B) in which nanoluciferase (NLuc) signal provides a readout of uORF2 repressiveness and firefly luciferase (FLuc) signal serves as normalization for transfection efficiency. We confirmed that uORF2 repressiveness depends on its translation and the terminal diproline-dependent elongating ribosome stall (Fig. 2C, top three rows compared to bottom wild-type). We used this *UL4*-based luciferase reporter to quantitatively dissect the kinetics of uORF-mediated translational regulation.

We complemented our experimental measurements with computational kinetic modeling of proposed models of uORF regulation (Fig. 1). We aimed to find unique modeling predictions that would allow us to experimentally distinguish between the different models of uORF regulation. We specified the kinetics of each of the proposed models of uORF regulation using PySB, a rule-based framework for compact model specification^52^. We then expanded the model into the BioNetGen modeling language syntax53 and inferred a reaction dependency graph for efficient simulation^39^. Next, we stochastically simulated the models using an agent-based Gillespie algorithm implemented in NFSim^54^. The molecules and reactions within the kinetic model are shown in Fig. S1A and Fig. S1B, respectively, and described in detail in the methods section. We experimentally tested predictions from this computational modeling and used the results to refine our model specifications. This iterative cycle of experimental testing and computational modeling constitutes our platform for differentiating between proposed uORF regulatory models.

While many rates of steps of translation have been estimated using single-molecule or ribosome profiling methods^55–58^, several critical parameters specific to *UL4* uORF2 have not been estimated. Therefore, we first calibrated our computational models to our reporter measurements on wild-type or mutant uORF2 (Fig. 2C) to derive estimates for these remaining, unknown parameters (marked with “This work” in Table 1). We used previously generated estimates for kinetic parameters not directly measured in our work (Table 1). We did not fit the constitutive repression and regulated re-initiation models (Fig. 1A,E) to our reporter measurements (Fig. 2C) since these models cannot account for the critical role of the *UL4* uORF2 elongating ribosome stall in regulating main ORF translation in single uORF transcripts.

**Table 1.** Modeling parameter ranges and reporter measurement (Fig. 2C) fit values.

| Parameter | Value range | Fit value (80S-hit dissociation) | Fit value (queuing-mediated enhanced repression) | Fit value (collision-mediated 40S dissociation) | Reference |
| --- | --- | --- | --- | --- | --- |
| $k_{cap\ bind}$ ($s^{-1}$) | 0.02-0.06 | 0.016 | 0.023 | 0.025 | This work<sup>56-58</sup> |
| $k_{scan}$ (residues/s) | 1-10 | 5 | 5 | 5 | <sup>38,99</sup> |
| $k_{start\ uORF2\ WT}$ ($s^{-1}$) | unknown | 0.1 | 0.5 | 0.5 | This work |
| WT uORF initiation (%) | unknown | 2 | 10 | 10 | This work |
| $k_{start\ uORF2\ strong\ Kozak}$ ($s^{-1}$) | unknown | 20 | 5 | 5 | This work |
| strong Kozak uORF initiation (%) | unknown | 80 | 50 | 50 | This work |
| $k_{elong}$ (residues/s) | 3-10 | 5 | 5 | 5 | <sup>55-58</sup> |
| $k_{elong\ stall}$ (residues/s) | 0.001 | 0.001 | 0.001 | 0.001 | <sup>101</sup> |
| $k_{terminate}$ ($s^{-1}$) | 0.5-5 | 1 | 1 | 1 | <sup>55</sup> |
| $k_{terminated\ ssu\ recycle\ uORF}$ ($s^{-1}$) | unknown | 2 | 5 | 5 | This work |
| Re-initiation (%) | unknown | 75 | 50 | 50 | This work |
| $k_{dissociate}$ ($s^{-1}$) | unknown | 2 | 0 | 2 | This work |
| uORF length (residues) | 21 | 21 | 21 | 21 | <sup>32</sup> |

Simulations of the queuing-mediated enhanced repression (Fig. 1C) and collision-mediated 40S dissociation (Fig. 1D) models readily recapitulate measurements on NLuc protein output in wild-type and mutant *UL4* reporters (Fig. 2D, triangles and squares). The 80S-hit dissociation model (Fig. 1B), modified to include an elongating ribosome stall within the uORF, also recapitulates the reporter measurements (Fig. 2D, circles). However, this modified 80S-hit dissociation model requires the difference between the stronger Kozak and wild-type uORF initiation to be quite large (80% vs. 2% compared to 50% vs. 10% for 2 other models mentioned above, Table 1). The derived ribosome loading rates (~0.02/s for all three of these models (Fig. 1B-D) are in line with literature estimates^56–58^. Although we do not measure how often terminating ribosomes at uORF2 re-initiate, the fractions derived here (50-70%, Table 1) are within the range of measured re-initiation fractions across mRNAs with different sequence features^40–43^. A complete description of the derivation of model parameters marked with a reference of “This work” can be found in the methods section. Given our computational recapitulation of experimental data, we then used our computational modeling platform to predict how translation would be perturbed following variations in other kinetic parameters.

### Computational modeling predicts that different models of uORF regulation have unique parameters important for buffering

While many kinetic parameters could be varied to help distinguish between proposed models of uORF regulation (Fig. 1), we honed in on the rate of ribosome loading onto the mRNA for two key reasons. Firstly, this rate is reduced endogenously in response to a variety of cellular and environmental signals. Amino acid deprivation, ribosome collisions, dsRNA viral infection, unfolded proteins, and heme deprivation are sensed by one of the four eIF2α kinases (GCN2, PKR, PERK, and HRI) to reduce the concentration of eIF2α-containing ternary complexes (TCs)^59–61^. A reduction in the concentration of eIF2α-containing TCs reduces the rate of ribosome loading. Viral infection also leads to reduced ribosome loading via interferon-induced proteins with tetratricopeptide repeats (IFITs)^62^. Cellular stress also reduces ribosome loading via inhibition of mTOR and sequestration of eIF4E by hypophosphorylated 4EBP^63^. Secondly, translated, repressive uORFs are enriched in transcripts buffered against reduced ribosome loading^64–67^. Therefore, we were particularly interested in varying this ribosome loading rate to investigate if and how uORFs provide this buffering across various proposed models. For each of the five surveyed models of uORF regulation (Fig. 1), we investigated what uORF parameters combinations, if any, allow buffering against reduced ribosome loading rates.

We use the term ‘buffer’ to describe the observation of main ORF translation decreasing less than expected, or even increasing, with reduced ribosome loading in comparison to the constitutive repression model (Fig. 1A). Importantly, buffering requires an interaction between ribosome loading and the degree of translational repression. We use buffering as an overarching term that encompasses both resistance and preferred translation. Resistance refers to main ORF translation reduced to a lower extent than the constitutive repression model when ribosome loading is reduced. Preferred translation refers to increased main ORF translation when ribosome loading is reduced. The constitutive repression model (Fig. 1A) has no buffering (Fig. 3A) since its repression is independent of the ribosome loading rate.

**Figure 3.**
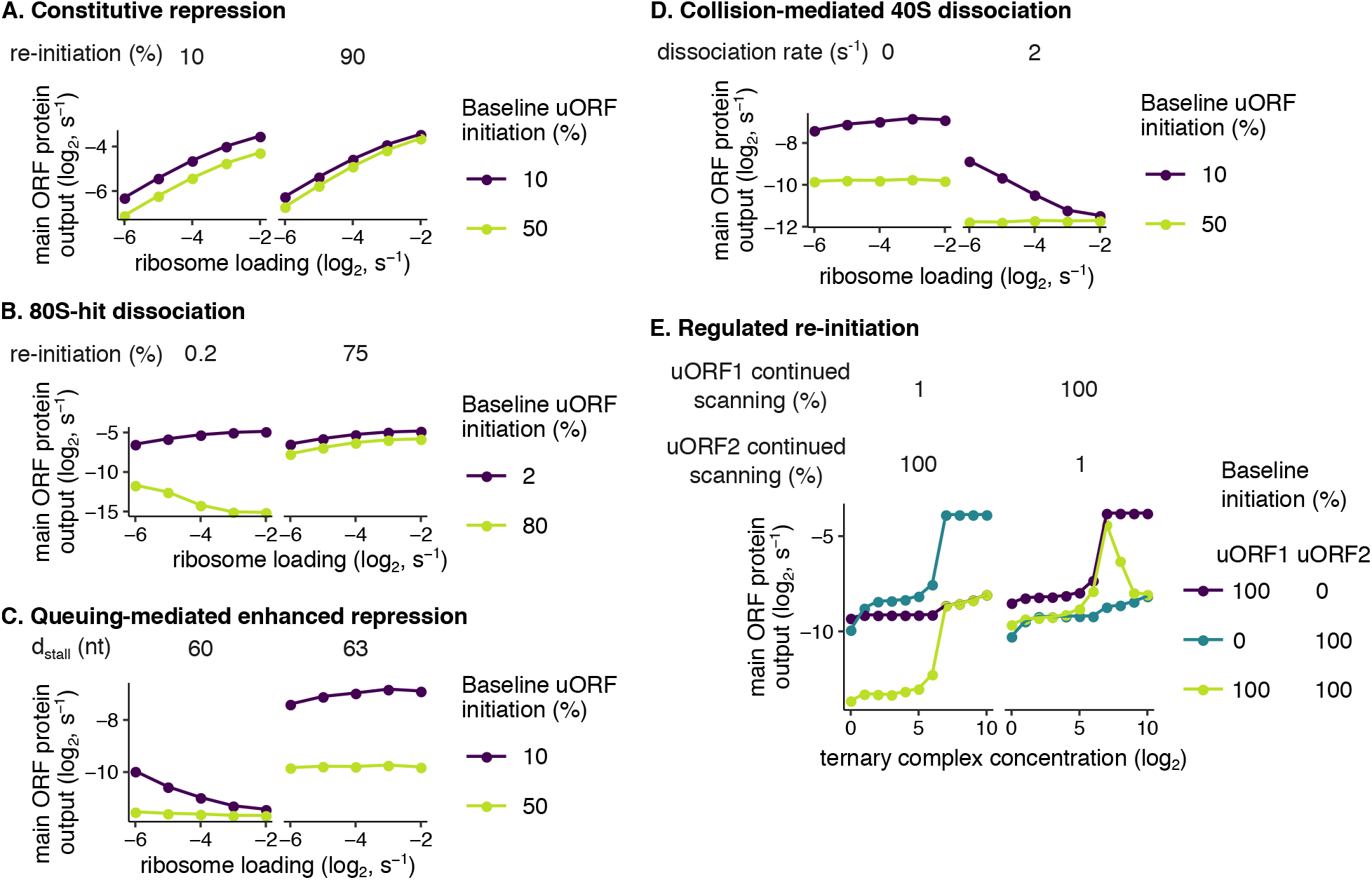
Kinetic modeling predicts translational buffering by uORFs. Buffering refers to a less than expected decrease (small positive slope), or even increase (negative slope), in main ORF translation with reduced ribosome loading. **(A)** The constitutive repression model, without an elongating ribosome stall, has no buffering behavior. In this model, buffering does not occur at any uORF initiation or re-initiation rates. uORFs simply siphon away scanning ribosomes from the main ORF. **(B)** Buffering in the 80S-hit dissociation model depends on uORF initiation and re-initiation frequencies^38^. For buffering to occur in this model, uORFs must initiate well enough to have elongating ribosomes hit 3’ scanning ribosomes (yellow-green line). uORFs must also not continue scanning at high frequencies following termination (left panel); frequent continuation of scanning coupled with high uORF initiation allows many scanning ribosomes to make it to the main ORF. The uORF is 100 residues long. Buffering occurs better for longer uORFs that have more time for elongating ribosomes to hit 3’ scanning ribosomes (Fig. S2A, yellow-green line). The dissociation rate is 200s^-1^, so 99% of scanning ribosomes hit by 5’ elongating ribosomes dissociate rather than continue scanning. The scanning rate is 2 nt/s, and the elongation rate is 2 codons/s. There is no elongating ribosome stall in this model. **(C)** Buffering in the queuing-mediated enhanced repression model depends on *d_stall_*. the distance between the uORF start codon and elongating ribosome stall. In this model, uORF initiation can increase above baseline with increased ribosome loading when the distance between the uORF start codon and elongating ribosome stall is an integer multiple of the ribosome footprint (30 nt, left panel). When this condition is met, buffering occurs. For *d_stall_* values of 60 and 63 nt, the uORF length is 21 and 22 codons, respectively. **(D)** Buffering in the collision-mediated 40S dissociation model depends on the dissociation rate. Here, *d_stall_* is 63 nt; with a low dissociation rate, this model reduces to the queuing-mediated enhanced repression model. **(E)** Buffering in the regulated re-initiation model depends on uORF initiation and continued scanning. For buffering to occur, several conditions must be met. At least 2 uORFs are required, both of which must be well-translated (yellow-green line). Continued scanning following termination at the first uORF must be frequent, and continued scanning following termination at the second downstream uORF must be rare (right panel). The second downstream uORF is 3 residues long. There is no elongating ribosome stall in this model. uORFs are located 25 nt from the 5’ cap. 99% of scanning ribosomes that make it to the main ORF will initiate translation; 1% will leaky scan. Unless otherwise stated, parameters (Table 1) obtained from calibrating models to reporter measurements on wild-type or mutant uORF2 (Fig. 2C) are used here. Ribosome loading is the *k_cap bind_* rate for non-regulated re-initiation models. We model changes in ribosome loading via changes in *k_cap bind_* as that rate is easier to match to *in vivo* estimates of ribosome loading. However, buffering in the regulated re-initiation model is dependent on an eIF2α phosphorylation mechanism; we instead vary the number of ternary complexes in this model. Error bars of simulated data are smaller than data points.

The 80S-hit dissociation model (Fig. 1B) displays buffering (Fig. 3B, left panel, yellow-green line) in agreement with previous work^38^. This behavior arises because the number of 5’ elongating ribosomes that collide with scanning ribosomes correlates with the ribosome loading rate. However, buffering depends on strongly initiating uORFs, minimal re-initiation, and longer uORFs (Fig. 3B, left panel, yellow-green line, Fig. S2A) as observed previously^38^. These observations can be rationalized as follows. Strong uORF initiation generates sufficient elongating ribosomes that hit and knock off 3’ scanning ribosomes. Minimal re-initiation prevents the many uORF-translating ribosomes from also translating the main ORF. Longer uORFs offer more time for elongating ribosomes to catch up, hit, and knock off 3’ scanning ribosomes. Nevertheless, most eukaryotic uORFs only weakly initiate translation and are short^1,3,4,8-11,31^. *UL4* uORF2 is 22 codons long, and we estimate reinitiation to be frequent (Table 1). Accordingly, buffering is no longer observed (Fig. S2B) in this model when parameters derived from control *UL4* variants (Table 1) are used.

The queuing-mediated enhanced repression model^23^ (Fig. 1C) displays buffering behavior (Fig. 3C, left panel, purple line) since the number of scanning ribosomes that initiate translation at the uORF is dependent on the rate of ribosome loading. In this model, reduced ribosome loading decreases the average queue length of ribosomes behind the elongation stall and thus, also the fraction of ribosomes that initiate at uORF2 (Fig. S2C, left panel). Unlike the 80S-hit dissociation model (Fig. 1B), weakly initiating uORFs, such as *UL4* uORF2, still confer buffering in this model (Fig. 3C, left panel, purple line).

In the queuing-mediated enhanced repression model, enhanced uORF initiation and, therefore, buffering are sensitive to the distance between the uORF start codon and the elongating ribosome stall (*d_stall_*) (Fig. S2C and Fig. 3C, left vs. right panels). This sensitivity arises because *d_stall_* determines if the P-site of a queued scanning ribosome is correctly positioned over the uORF start codon to productively increase uORF initiation (Fig. S2C, left panel). In the idealized case of homogeneously sized ribosomes (30 nt footprints^55,68^) and strict 5’-3’ scanning, *d_stall_* must be an integer multiple of 30 nt for buffering to occur. This strong dependence of buffering on *d_stall_* is relaxed when backward scanning^41,69-71^ occurs with a high rate (Fig. S2D, middle panel). However, even a high rate of backward scanning is insufficient to compensate for misalignment between the uORF start codon and the P-site of the queued scanning ribosome (Fig. S2D, middle panel). To simplify our modeling interpretation, we considered *UL4* uORF2, that is 22 codons long, to be 21 codons so that a queue behind the terminating ribosome stall positions a scanning ribosome’s P-site exactly on the start codon.

The collision-mediated 40S dissociation model (Fig. 1D) displays buffering (Fig. 3D, right panel, purple line) because the number of scanning ribosomes that collide with 3’ stalled elongating ribosomes is dependent upon the rate of ribosome loading. Buffering in this model requires the collision-induced 40S dissociation rate to be somewhat fast (Fig. 3D, right vs. left panels, Fig. S2E, teal and yellow-green lines). If this rate is too low (for example, 0 in Fig. 3D, left panel), this model reduces to the queuing-mediated enhanced repression model (Fig. 1C). However, unlike the queuing model, the collision-mediated 40S dissociation model is not sensitive to the distance between the stall and the start codon (Fig. S2F, purple lines). As in the queuing model (Fig. 1C), weakly initiating uORFs, such as *UL4* uORF2, can still confer buffering (Fig. 3D, right panel, purple line) in the collision-mediated 40S dissociation model (Fig. 1D). This effect arises because, unlike in the 80S-hit dissociation model (Fig. 1B), the elongation stall is now rate limiting for main ORF translation. Therefore, the mere presence of an elongation stall within the queuing-mediated enhanced repression and collision-mediated 40S dissociation models imparts some degree of buffering.

In the regulated re-initiation model (Fig. 1E), buffering is observed (Fig. 3E, right panel, yellow-green line) because termination at the first uORF is followed by reinitiation at either the second downstream uORF or the main ORF depending on the ternary complex concentration. Buffering in the regulated re-initiation model (Fig. 1E) depends upon the initiation efficiency and continued scanning fractions of the two uORFs (Fig. 3E). Continued scanning following termination at the first uORF must be frequent while continued scanning following termination at the second downstream uORF must be rare. Higher ternary complex concentration biases towards initiation at the second downstream uORF (Fig. S3A). Reductions from high ternary complex concentrations biases towards main ORF initiation; therefore main ORF translation can increase with decreased ribosome loading.

As such, our computational results provide the first systematic comparison of different mechanisms of uORF-mediated regulation (Fig. 1) and enable their comparison with experimental measurements below.

### *UL4* uORF2 buffers against reductions in main ORF translation from reduced ribosome loading in an elongating ribosome stall-dependent manner

We next tested whether the computational predictions of uORF-mediated buffering (Fig. 3) can be observed experimentally with *UL4* uORF2. To this end, we experimentally varied the rate of ribosome loading and measured effects on main ORF translation using our reporter system (Fig. 2B). Since no-stall uORF2 variants have similar protein expression to the no-start uORF2 variants (Fig. 2C), luciferase signal from the no-stall uORF2 variants provides a readout of the ribosome loading rate. If buffering were absent, then we would expect NLuc translation to be reduced equally between no-stall and wild-type variants when ribosome loading is reduced.

We used three strategies to vary the rate of ribosome loading. We first used stem loops near the 5’ cap that reduce the rate of 43S-cap binding without affecting mRNA stability (Fig. 4A)^72,73^. We varied the degree to which ribosome loading is reduced by altering the GC content of the stem loops. We observe that NLuc translation decreases less with reduced ribosome loading for the wild-type *UL4* reporter in comparison to the no-stall UL4 variant (Fig. 4A, left panel, yellow vs. gray circles). Therefore, wild-type uORF2 imparts resistance to reductions in NLuc translation from stem loop-mediated reduced ribosome loading. When the wild-type data are normalized by the no-stall data, NLuc translation negatively correlates with ribosome loading, indicative of buffering against reduced ribosome loading by wild-type uORF2 (Fig. 4A, right panel).

**Figure 4.**
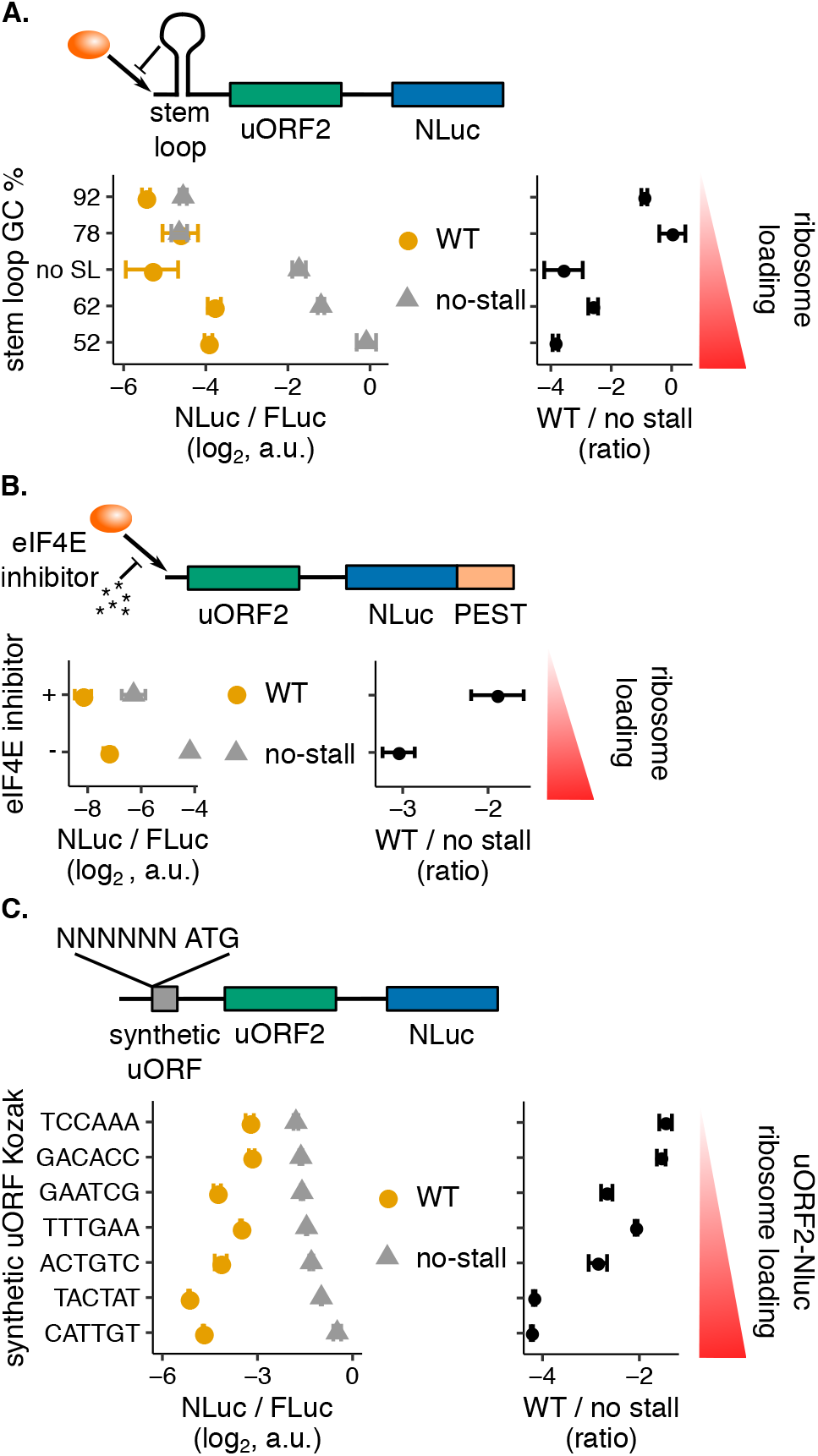
The human cytomegaloviral uORF2 buffers against reductions in main ORF translation. The human cytomegaloviral *UL4* uORF2 is used in the dual-luciferase assay (Fig. 2B) in conjunction with various mechanisms to reduce ribosome loading. **(A)** Ribosome loading is reduced using stem loops^73^ with the indicated GC percentages. The −30 kcal/mol stem loops are positioned 8 bp from the 5’ cap. The no-stem loop data has a 5’ UTR CAA repeat instead of a stem loop. The 5’ UTR is 287 nt long. Data are normalized to a no-uORF start codon control without a stem loop. **(B)** Ribosome loading is reduced using the drug 4E1RCat (116 *μ*M)^74^ that disrupts the interaction between eIF4E and eIF4G. NanoLuc has a C-terminal PEST tag to increase protein turnover^75^ for the 3-hour drug treatment. The 5’ UTR is 236 nt long. Data are normalized to a no-uORF start codon control without a PEST tag. **(C)** Ribosome loading onto the uORF2-NanoLuc portion of the transcript is reduced using a 5’ synthetic uORF: ATG GGG TAG. The synthetic uORF Kozak is varied to alter ribosome loading. The data are vertically ordered by the no-stall means. The 5’ UTR is 262 nt long. Data are normalized to a no-uORF start codon control without a synthetic uORF. Right panels in A,B, C show wild-type (WT) mean values normalized by the corresponding no-stall values. The no-stall uORF2 mutants lack their terminal diproline motifs (P22A mutation). Error bars show standard error of mean NLuc / FLuc ratios over 3 technical replicates.

We also reduced ribosome loading with the drug 4E1RCat, which disrupts the interaction between the 5’ cap-binding eIF4E and the scaffold eIF4G (Fig. 4B)^74^. We added a PEST tag75 to increase the turnover of the NLuc protein to more accurately measure changes in main ORF translation during drug treatment. NLuc translation of wild-type *UL4* reporter decreases less in comparison to the no-stall control upon 4E1RCat treatment (Fig. 4B, left panel, yellow vs. gray circles), indicative of resistance. Again, when the wild-type data are normalized by the no-stall data, we observe that NLuc translation negatively correlates with ribosome loading, indicative of general buffering against reduced ribosome loading by wild-type uORF2 (Fig. 4B, right panel).

Finally, we added a short, synthetic uORF, 5’ to the *UL4* uORF2, that siphons scanning ribosomes away from uORF2 (Fig. 4C). We varied the degree of ribosome siphoning by varying the Kozak context of the synthetic uORF, which in turn determines the rate of ribosome loading onto the uORF2-NLuc portion of the mRNA. Here, we observe that more NLuc is produced in the wild-type *UL4* reporter as scanning ribosomes are increasingly siphoned off by improving the Kozak context of the synthetic uORF (Fig. 4C, left panel, yellow circles). While resistance is observed with the other strategies of reduced ribosome loading (Fig. 4A-B, left panels), preferred translation is observed here (Fig. 4C, left panel), perhaps because ribosome loading is reduced only within the transcript rather than at the 5’ cap. However, similar to the other two strategies, when the wild-type data are normalized by the no-stall data, NLuc translation negatively correlates with ribosome loading, indicative of general buffering against reduced ribosome loading by wild-type uORF2 (Fig. 4C, right panel).

### Distance between the start codon and stall does not systematically regulate uORF repressiveness or buffering

Given our experimental data of uORF2-mediated buffering of *UL4* reporters (Fig. 4), we narrowed our focus from the five surveyed models (Fig. 1) to the two (Fig. 1C-D) most relevant for *UL4* uORF2: the queuing-mediated enhanced repression (Fig. 1C) and collision-mediated 40S dissociation models (Fig. 1D). These two models are computationally predicted to confer buffering in an elongating ribosome stall-dependent manner without needing multiple uORFs (Fig. 3C-D). To differentiate between these models, we turned to our computational modeling prediction that, only in the queuing-mediated enhanced repression model (Fig. 1C), main ORF protein output is sensitive to the distance between the uORF start codon and elongating ribosome stall (Fig. 3C). Our computational modeling of the queuing-mediated enhanced repression model (Fig. 5A, yellowgreen line) predicts two broadly spaced clusters of main ORF protein output. Protein output from the main ORF is repressed when the start codon-stall distance is an integer multiple of the ribosome size. Protein output from the main ORF is high when the start codon-stall distance is not an integer multiple of the ribosome size. In contrast, the collision-mediated 40S dissociation model (Fig. 1D) predicts a much lower effect of *d_stall_* on uORF repressiveness (Fig. 5A, left panel, purple line). The residual effect of *d_stall_* on uORF repressiveness (Fig. 5A, left panel, purple line) in the collision-mediated 40S dissociation model (Fig. 1D) arises because the dissociation rate is low enough to allow rare queuing.

**Figure 5.**
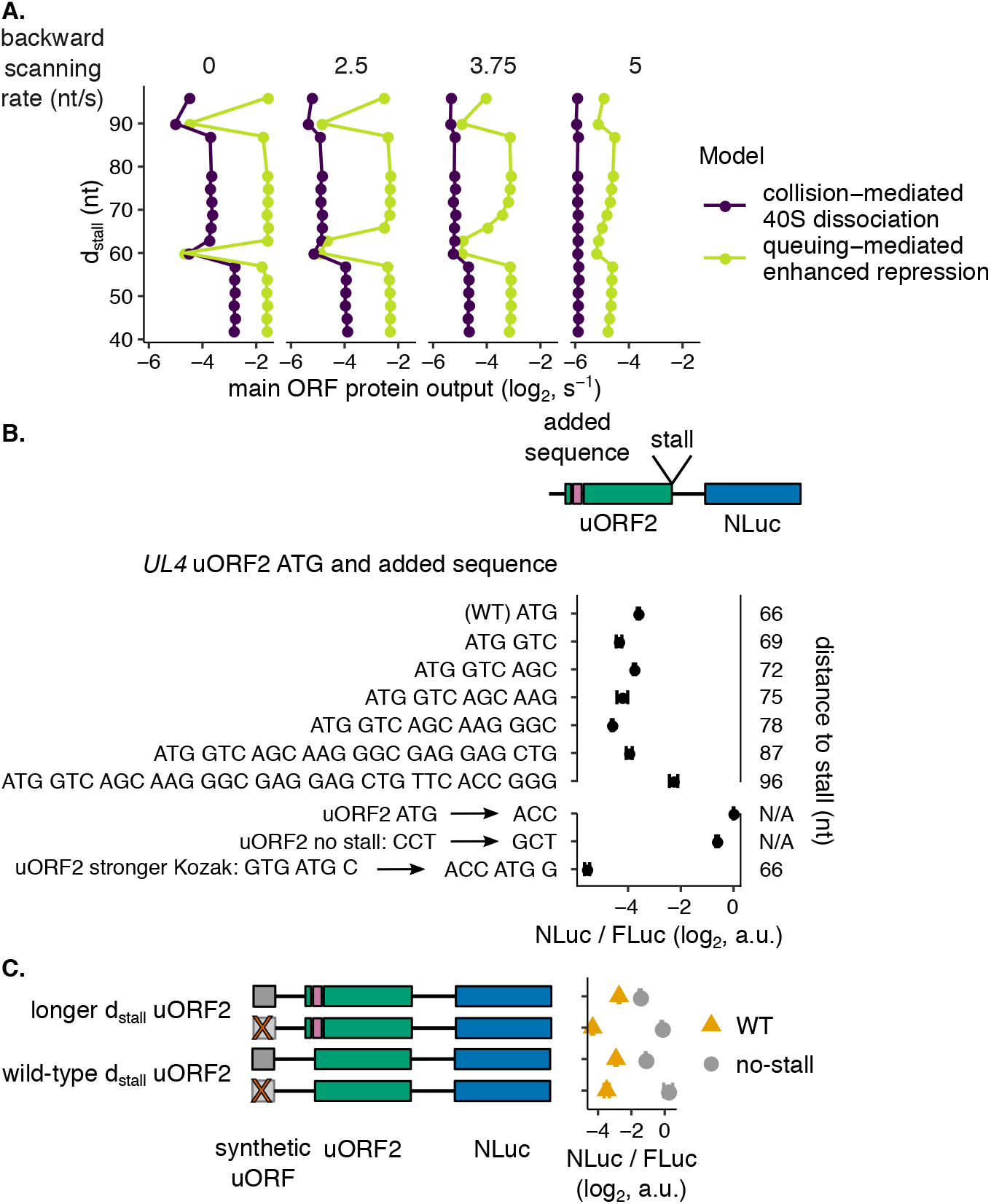
Changes to the distance between the human cytomegaloviral uORF2 start codon and elongating ribosome stall do not change repressiveness or buffering as predicted by computational modeling of the collision-mediated 40S dissociation model. **(A)** Computational modeling predicts greater changes in uORF repressiveness with changes in *d_stall_* in the queuing-mediated enhanced repression model. Fast backward scanning abolishes this periodicity in main ORF translation across varied *d_stall_*. *d_stall_* refers to the distance between the start codon and elongating ribosome stall. As backward scanning increases in rate (moving right along panels), the collision-mediated enhanced repression model loses periodicity (middle panel, purple line) before the queuing-mediated enhanced repression model (right panel, yellow-green line). Parameters that best recapitulated reporter measurements on wild-type or mutant uORF2 (Fig. 2C, Table 1) are used here. The forward scanning rate is 5 nt/s. Data are normalized to a no-uORF start codon control. Error bars of simulated data are smaller than data points. **(B)** Experimentally varying the distance between the human cytomegaloviral uORF2 start codon and elongating ribosome stall does not systematically affect its repression of main ORF translation. The human cytomegaloviral *UL4* uORF2 is used in the dual-luciferase assay (Fig. 2B) in conjunction with various length inserts from the N-terminus of the *EYFP* main ORF. The *EYFP* main ORF sequence is inserted directly 3’ to the uORF2 start codon. The added sequence increases the distance between the uORF2 start codon and elongating ribosome stall. The bottom three controls improve the uORF2 Kozak context, remove the start codon, and remove the elongating ribosome stall. Error bars show standard error of mean NLuc / FLuc ratios over 3 technical replicates. Data are normalized to a no-uORF start codon control. **(C)** Experimentally varying the human cytomegaloviral uORF2 *d_stall_* does not strongly regulate the capacity of buffering against reductions in main ORF translation. Ribosome loading is reduced with a 5’ synthetic uORF: ATG GGG TAG. The no-stall uORF2 mutants lack their terminal diproline motifs (P22A mutation). No synthetic uORF mutants (ATG to AAG) are depicted by transparent, gray bars with red Xs and have a higher relative ribosome loading rate onto the uORF2-NLuc portion of the transcript. The distance between the uORF2 start codon and elongating ribosome stall is varied as indicated by adding 6 nt, GTC AGC, from the N-terminus of the *EYFP* main ORF. Data are normalized to a no-uORF start codon control without a synthetic uORF.

Backward scanning is predicted to diminish the periodicity in main ORF translation with varying *d_stall_* lengths in both models (Fig. 5A). However, backward scanning occurring as fast as forward scanning (~ 5 nt/s) is required to abolish the periodicity in the queuing model (Fig. 5A, right panel, yellow-green line). While there are estimates of how far ribosomes can backward scan^69–71^, we are not aware of any backward scanning rate estimates. It is unlikely that the rate of backward scanning approaches the rate of forward scanning (5 nt/s here) given the 5’-3’ directionality of scanning. Slower backward scanning (~ 3.75 nt/s) is sufficient to abolish periodicity in the collision-mediated 40S dissociation model (Fig. 5A, middle panel, purple line). This effect is not surprising given that the presence of periodicity in the latter model arises from rare queuing behavior. Therefore, our computational predictions of greater periodicity in main ORF translation across varied *d_stall_* in the queuing model hold even with backward scanning.

We then experimentally varied the distance between the start codon and stall of *UL4* uORF by adding codons to the 5’ end of uORF2. We observe less than 2-fold changes in translational regulation (Fig. 5B, top 7 rows) with no systematic trend with variations in uORF2 length, which is inconsistent with computational modeling predictions of the queuing-mediated enhanced repression model (Fig. 5A, left panel). Our longest uORF mutant here (96 nt from start to stall in Fig. 5B) is less repressive, but this effect may be due to decreased elongating ribosome stall strength. In this case, the 32 amino acid-long nascent peptide can extend out of the exit tunnel and be bound by additional factors^27,28^. Thus, our experimental data does not match computational predictions of main ORF translation sensitivity to *d_stall_* in the queuing-mediated enhanced repression model (Fig. 1C), and better supports the collision-mediated 40S dissociation model (Fig. 1D).

In the queuing-mediated enhanced repression model (Fig. 1C), buffering is uniquely predicted to be sensitive to the distance between the uORF start codon and elongating ribosome stall (Fig. 3C). We, therefore, asked whether or not buffering would still be experimentally observed with a disruption in this distance. Using our synthetic uORF method of reducing ribosome loading (Fig. 4C), we observe that a 6 nt longer *d_stall_* uORF still buffers against reduced ribosome loading (Fig. 5C, top two rows compared to bottom two rows). Since backward scanning of 15-17 nt has been observed^69–71^, one would expect that buffering would still be predicted in the queuing model even with an increase in *d_stall_* of 6 nt. However, our computational modeling predicts that even very fast backward scanning does not restore buffering when *d_stall_* is disrupted by 6 nt (Fig. S2D, right panel). Thus, our experimental data does not match computational predictions of buffering sensitivity to *d_stall_* in the queuing-mediated enhanced repression model (Fig. 1C), but is consistent with the collision-mediated 40S dissociation model (Fig. 1D).

### Several human uORFs have repressive terminal diproline motifs

Given that the elongating ribosome stall in the human cytomegaloviral *UL4* uORF2 is dependent on a terminal diproline motif, we asked whether there are other human uORFs similarly ending in diproline motifs that are also repressive. We searched for such uORFs in three databases: a comprehensive database of ORFs in induced pluripotent stem cells and human foreskin fibroblasts with 1,517 uORFs^6^, a database integrated from *de novo* transcriptome assembly and ribosome profiling with 3,577 uORFs^76^, and a database of proteins less than 100 residues in size derived from literature mining, ribosome profiling, and mass spectrometry with 1,080 uORFs^77^. We identified several human transcripts with uORFs with terminal diproline motifs: *C1orf43, C15orf59, TOR1AIP1*, and *ABCB9* (Fig. 6). We replaced *UL4* uORF2 in our reporter (Fig. 2B) with these human uORFs. We created terminal proline to alanine and no-start-codon mutants and measured the effects of these mutations on NLuc expression relative to the wild-type human uORFs. While many of the tested uORFs are repressive (Fig. 6, yellow vs. blue), unlike the human cytomegaloviral uORF2, these human uORFs still generally repress NLuc expression without their terminal diproline motif (Fig. 6, gray vs. blue), indicating additional contributions to translational repression from siphoning of scanning ribosomes at the start codon and other residues in the nascent peptide.

**Figure 6.**
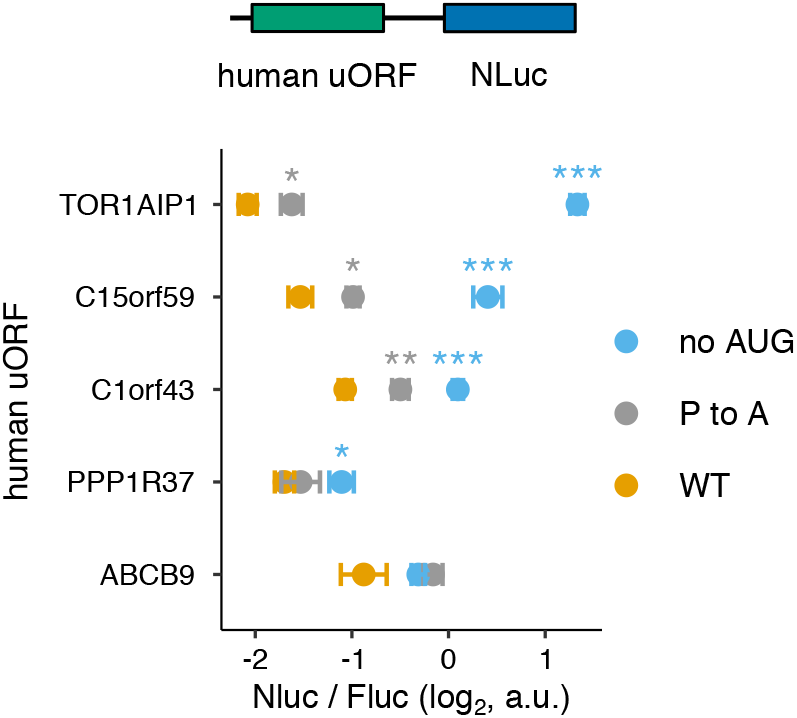
Several human uORFs have repressive terminal diproline motifs. Terminal diproline motif-containing human uORFs are used in the dual-luciferase assay (Fig. 2B). The terminal proline codon in each uORF is mutated to an alanine codon in the P to A mutant. Start codons are mutated to ACC for the no-AUG mutants. P values comparing the indicated mutants to the wild-type are shown after running a two sample t-test: N.S. (P > 0.05), * (0.01 < P < 0.05), ** (0.001 < P < 0.01), **\*** (P < 0.001). All comparisons with no shown significances are N.S. Data are normalized to a no-*UL4*-uORF2 start codon control.

## Discussion

In this study, we use a combination of computational modeling and experimental reporter measurements to dissect the kinetics of uORF-mediated translational regulation of the *UL4* mRNA of human cytomegalovirus. We find that elongating ribosome stalls in *UL4* uORF2 buffer against reductions in main translation due to reduced ribosome loading (Fig. 4). Using an experimentally-integrated modeling approach, we differentiate between models of regulation that can explain this observation. Our computational framework allows easy specification and efficient simulation of several previously proposed kinetic models of uORF regulation (Fig. 1). Thus, we can predict which models of uORF regulation of translation allow buffering and which parameters are key for buffering in each model (Fig. 3). While uORFs are enriched in stress-resistant transcripts, not all uORFs provide buffering^64^. To our knowledge, our work is the first systematic investigation of what uORF metrics impart buffering in each kinetic model of uORF regulation.

uORFs are generally thought to simply siphon away scanning ribosomes from main ORFs, but this simple behavior in the constitutive repression model (Fig. 1A) is not predicted to provide buffering (Fig. 3C)^64–67^. Instead, we find that 5’ UTRs containing one (or some combination) of the following enable buffering main ORF translation: scanning ribosome dissociation due to 80S hits from the 5’ (Fig. 1B), a single uORF with an elongating ribosome stall (Fig. 1C-D), or multiple uORFs acting through the regulated re-initiation model (Fig. 1E).

Long, well-initiating uORFs that do not re-initiate well allow buffering (Fig. 3B, left panel, yellow-green line, Fig. S2A, yellow-green line) in the 80S-hit model (Fig. 1B), but these requirements are at odds with the typically short and poorly initiating nature of known uORFs^1,3,4,8–11^. Consequently, when we use parameters specific to *UL4* uORF2 for the 80S-hit model (Table 1), namely that uORF2 initiates poorly, re-initiates well, and is not very long, buffering is no longer predicted (Fig. S2B).

Our modeling of the regulated re-initiation model (Fig. 1E) agrees (Fig. 3E and Fig. S3A) with previous work^35,36^ that buffering requires 1) two well-translated uORFs, and 2) frequent and rare continued scanning aftertermination at uORFs 1 and 2, respectively. The 30% of human transcripts that contain multiple uORFs can display buffering by the regulated re-initiation model; however, about 25% of human transcripts only have one uORF2 and are unable to provide buffering via this model.

We narrowed our focus to the two models (Fig. 1CD) that are most pertinent to *UL4* uORF2. Both the queuing-mediated enhanced repression (Fig. 1C) and collision-mediated 40S dissociation (Fig. 1D) models are predicted (Fig. 3C-D) to allow buffering with weakly initiating uORFs and elongating ribosome stalls. Both of these models include an elongating ribosome stall and only require a single uORF for buffering (Fig. 3C-D). Computational modeling not only predicted this buffering behavior but also allows us to differentiate between these two models. We predict that the queuing-mediated enhanced repression model (Fig. 1C) is uniquely sensitive to the distance between the uORF start codon and elongating ribosome stall (Fig. 5A, yellow-green line, Fig. 3C, purple lines, Fig. S2F, purple lines). We experimentally vary this distance and do not find any systematic changes to either main ORF protein output (Fig. 5B) or buffering (Fig. 5C). Based on our results, we propose that scanning ribosomes dissociate rather than queue when encountering a 3’ stalled elongating ribosome on uORF2 of *UL4* mRNA.

While we have no direct evidence of scanning ribosome dissociation, the requirement of crosslinking to retain scanning ribosomes during ribo-seq preparations^78,79^ hints at their labile nature and potential dissociation *in vivo*. This dissociation could serve to maintain the free pool of 40S ribosomal subunits while still allowing regulation of main ORF translation. Collisions between scanning ribosomes and their subsequent dissociation have also recently been proposed in a model of initiation RQC^80^. Collisions between scanning and elongating ribosomes and subsequent quality control are not well understood; What we describe as scanning ribosome dissociation here may be rescue by a quality control pathway.

Although our data from *UL4* uORF2 does not support the queuing-mediated enhanced repression model (Fig. 1C)^23^, this model might be relevant in other scenarios. Translation from non-cognate start codons is resistant to cycloheximide, perhaps due to queuing-mediated enhanced initiation, but sensitive to reductions in ribosome loading^81^. Loss of eIF5A, which helps paused elongating ribosomes continue elongation, increases 5’ UTR translation in 10% of studied genes in human cells, augmented by downstream in-frame pause sites within 67 codons, perhaps also through queuing-mediated enhanced initiation^82^. There is also evidence of queuing-enhanced uORF initiation in the 23 nt long *Neurospora crassa* arginine attenuator peptide83 as well as in transcripts with secondary structure near and 3’ to start codons^84^. Additional sequence elements in the mRNA might determine whether scanning ribosome collisions result in queuing or dissociation.

*UL4* uORF2 is not unique in containing an elongating ribosome stall^22,23,26,47-51^. There are a variety of residues that may reduce the rate of elongation, either through changes in the activity of the peptidyl transferase, tRNA availabilities, or interactions between the nascent peptide and the ribosome^85,86^. uORFs are often short3 and may therefore be better poised to stall ribosomes; the nascent peptides may be less likely to be released due to interactions with the ribosome. Thus, a key role for elongation ribosome stalls in uORFs might be to enable buffering. While very few uORFs have been mechanistically characterized^1^, other elongating ribosome stall-containing uORFs, such as the peptide sequence sensitive uORFs that regulate human methionine synthase^47^ and antizyme inhibitor I^23^, might enable buffering. Conversely, uORFs in several single uORF transcripts known to buffer against stress, such as SLC35A4, C19orf48, and IFRD1^64^, might act through elongating ribosome stalls.

The computational models considered here can be readily extended to incorporate more complex mechanisms of translational control. For example, in our models, initiation proceeds via a cap-severed mechanism in which multiple scanning ribosomes can be present in the 5’ UTR at the same time. If we were to model cap-tethered initiation, strong uORF elongating ribosome stalls would eventually sever this connection, similar to how the cap-eIF-ribosome connection is severed during the usually longer translation of main ORFs^87–89^. It will be also interesting to consider the effect of cellular stress-reduced elongation rates90 and increased re-initiation^91^, both of which might regulate uORF-mediated buffering, as well as elongating ribosome dissociation through known quality control pathways^39,80,92-96^.

## Author Contributions

T.A.B. designed research, performed computations and experiments, analyzed data, and wrote the manuscript. A.P.G. conceived the project, designed research, and wrote the manuscript. A.R.S. conceived the project, designed research, performed computations and experiments, analyzed data, wrote the manuscript, supervised the project, and acquired funding.

## Acknowledgements

We thank members of the Subramaniam lab, the Basic Sciences Division, and the Computational Biology Program at Fred Hutch for assistance with the project and discussions and feedback on the manuscript. The computations described here were performed on the Fred Hutch Cancer Research Center computing cluster. This research was funded by NIH R21 AI156152 received by APG and NIH R35 GM119835, NSF MCB 1846521, and the Sidney Kimmel Scholarship received by ARS. This material is based upon work supported by the National Science Foundation Graduate Research Fellowship Program under Grant Numbers NSF DGE-1762114 and DGE-2140004. Any opinions, findings, and conclusions or recommendations expressed in this material are those of the author(s) and do not necessarily reflect the views of the National Science Foundation. This research was supported by the Genomics Shared Resource of the Fred Hutch/University of Washington Cancer Consortium (P30 CA015704) and Fred Hutch Scientific Computing (NIH grants S10-OD-020069 and S10-OD-028685). The funders had no role in study design, data collection and analysis, decision to publish, or preparation of the manuscript.

## Data and Code Availability

All experimental data and programming code associated with this manuscript is available at https://github.com/rasilab/bottorff_2022.

## Materials and Methods

### Plasmid construction

The parent cloning vector was created as follows. A commercial vector (Promega pGL3) with ampicillin resistance was used to clone nanoluciferase and firefly luciferase. NanoLuc expression is driven by a CMV promoter. Firefly luciferase expression is driven in the opposite direction within the plasmid and serves as an internal transfection control. The human cytomegaloviral *UL4* 5’ UTR was PCR amplified from a construct gifted from HCMV genomic DNA. To create mutant 5’ UTR versions of the parent pGL3-FLuc-NLuc vector, the vector was digested with KpnI/EcoRI unless otherwise noted. 1 or 2 PCR-amplified fragments with 20-30 bp homology arms were then cloned using isothermal assembly^97^. The stem loop73 5’ UTR mutants were cloned as follows. The stem loops were ordered as oligonucleotides with overhangs for ligation into ClaI and NotI sites. The oligonucleotides were annealed and used in PCR reactions to add CMV homology arms. An AAVS1 parent vector was digested with ClaI and NotI. These stem loops were then inserted into the ClaI/NotI restriction digested parent vector by isothermal assembly^97^. The stem loops were then PCR amplified off of this plasmid and inserted into the pGL3-Fluc-*UL4*-5’-UTR-NLuc parent vector described above. The several tested human uORFs were PCR amplified from human genomic DNA and inserted into a PstI/EcoRI digested parent. The inserted sequences were confirmed by Sanger sequencing. Kozak context and stall codon mutations were introduced in the PCR primers used for amplifying inserts before isothermal assembly. Standard molecular biology procedures were used for all other plasmid cloning steps^98^. Table S1 lists the plasmids described in this study. Key plasmid maps are available at https://github.com/rasilab/bottorff_2022 as SnapGene `.dna` files. Plasmids will be sent upon request.

### Cell culture

HEK293T cells were cultured in Dulbecco’s modified Eagle medium (DMEM 1X, with 4.5 g/L D-glucose, + L-glutamine, - sodium pyruvate, Gibco 11965-092) and passaged using 0.25% trypsin in EDTA (Gibco 25200-056).

### Dual-luciferase reporter assay

Plasmid constructs were PEI or Lipofectamine 3000 (Invitrogen, L3000-008) transiently transfected into HEK293T cells for 12-16h in 96, 24, or 12 well plates. If the plasmids were not transfected into a 96 well plate, then the cells were resuspended in 100 *μ*L media. Then, 20 *μ*L of these resuspended cells were added per well to a 96 well plate for the dual-luciferase assay. If the transfection was already in a 96 well plate, the ~110 *μ*L media was removed and replaced with 20 *μ*L media per well. The Promega dual-luciferase kit was used. Cells were lysed with 20 *μ*L ONE-Glo EX Luciferase Assay Reagent per well for three minutes to measure firefly (*Photinus pyralis*) luciferase activity. Then, 20 *μ*L NanoDLR Stop & Glo Reagent was added per well for 10 minutes to quench the firefly luciferase signal and provide the furimazine substrate needed to measure NanoLuc luciferase activity. Firefly luciferase activity serves as an internal control for transfection efficiency, and NanoLuc activity provides a readout of 5’ UTR regulation of NanoLuc translation.

### Kinetic modeling

We specify our kinetic models using the PySB interface52 to the BioNetGen modeling language53 (Fig. S1). The Python script is parsed by BioNetGen into a `.bngl` file and converted into an `xml` file for use as input to the agent-based stochastic simulator NFsim^54^.

### Molecules

Our kinetic models of eukaryotic translational control describe the interactions between 2 molecule types: mRNA and ribosome (composed of separate large and small subunits). Here, we describe these molecules’ components, states, and binding partners (Fig. S1A). mRNA molecules have the following components: 5’ end and codon sites (*c_i_*). The mRNA 5’ end can either be free of or occupied with a ribosome. The mRNA 5’ end must be free for a 43S to bind, which leaves the 5’ end blocked until the ribosome scans (or elongates) sufficiently 3’ downstream. The mRNA codon sites serve as bonding sites for the ribosome A site. Ribosomes, both scanning and elongating, have the following components: A site (*a*), 5’ side (*t* for trailing), and 3’ side (*l* for leading). These sites serve as bonding sites for either the mRNA (A site) or other ribosomes during collisions (5’ or 3’ side). Both scanning and elongating ribosomes have mRNA footprints of 10 codons in our simulations based on mammalian ribosome profiling data^55,68^.

### Reactions

We describe here each type of kinetic reaction in our models of eukaryotic translational control (Fig. S1B). We use a syntax similar to that of BioNetGen53 to illustrate the kinetic reactions. We scale ternary complex (TC, 100) and ribosome subunit numbers (100 each) to the single mRNA present in the simulation. Simulation of a single mRNA is sufficient to infer translation dynamics.

#### Initiation: PIC (43S) formation

Small ribosomal subunits must bind TCs to form preinitiation complexes (PICs, 43Ss) before loading onto mRNAs. We assume that PIC formation is irreversible. PIC formation is not rate-limiting in our simulations; we set the rate of 43S-cap binding (*k_cap bind_*) to be rate-limiting and to a total rate (independent of [43S]) to match the overall initiation rate to that of cellular estimates. Therefore, we arbitrarily set the second-order PIC formation rate (40S-TC binding rate, *k_ssu tc bind_*) to 0.01 * TC^-1^ * SSU^-1^ such that 100 40S-TC binding events occur per second, which is much higher than the 43S-cap binding rate.

#### Initiation: PIC (43S) loading onto mRNA

We model ribosome footprints at 30 nt following mammalian ribosome profiling data^55,68^. Therefore, PIC loading can occur when the 5’ most 30 nucleotides (nt) of the mRNA are not bound to any ribosomes. The rate at which PICs load onto the 5’end of the mRNA, *k_cap bind_*, is varied over a 100-fold range from the maximum ribosome loading rate, 0.125/s, based on single-molecule estimations in human cells^56^. PICs can load onto the mRNA when a ribosome footprint-sized region at the 5’ mRNA end is free of ribosomes. PIC loading results in the 5’ end being blocked until this ribosome scans or elongates past a ribosome footprint from the 5’ cap. We assume that PIC loading is irreversible.

#### Initiation: Scanning and start codon selection

The scanning rate is 5 nt/s following an estimate in a mammalian cell-free translation system^99^, and previous computational study^38^. Small ribosomal subunit A sites must be positioned exactly over start codons to initiate translation. The uORF start codon is 25 nt from the 5’ cap. We vary the rate at which this start codon selection occurs at the uORF in our modeling. Start codon selection releases the TC bound to the small ribosomal subunit. We assume that TC is regenerated instantaneously. The start codon selection rate divided by the sum of this start codon selection rate, the scanning rate, and the backward scanning rate equals the baseline initiating fraction. This calculation of the baseline initiating fraction will underestimate the initiating fraction in the case of correctly positioned 3’ ribosome queues (as in the queuing-mediated enhanced repression model). We assume that start codon selection is irreversible.

#### Elongation

Elongation results in the ribosome A site moving from codon *c_i_* to codon *c_i+1_*. The rate of elongation is set to 5 codons/s following single-molecule method and ribosome profiling estimates in mammalian cells of 3-18 codons/s^55–58,100^. Elongation may only proceed if there is no occluding 3’ ribosome; in other words, elongation may only proceed from codon *c_i_* to codon *c_i+1_* if the next 3’ ribosome’s A site is bound to a codon no more 5’ than *c_i+11_*. The elongation rate at the stall within the uORF is set to 0.001/s based on toeprinting assays over time with various inhibitory drugs^101^.

#### Termination, continued scanning, and re-initiation

Termination results in the dissociation of the large ribosomal subunit, but the small ribosomal subunit may continue scanning and subsequently re-initiate if a new TC is acquired before the next start codon is encountered. The termination rate is set to 1/s given that ribosome density tends to be higher at stop codons than within ORFs^55,102^. The recycling rate of terminated small ribosomal subunits after uORF translation is varied to model the effect of varied continued scanning after uORFs on the regulation of main ORF translation. The scanning rate divided by the sum of the scanning rate and this recycling rate equals the continued scanning fraction.

#### Collisions and dissociations

A collision between two ribosomes requires them to be separated by exactly one ribosome footprint in distance on the mRNA and result in bonds between the 5’ side of the leading (3’ most) ribosome and the 3’ side of the trailing (5’ most) ribosome. Abortive (premature) termination of ribosomes results in their dissociation from the mRNA and any collided ribosomes they are bound to. Different models have different non-zero dissociation rates. For instance in the 80S-hit model, the following rates are equal and non-zero: *k_scan term 5 hit 80s_*, *k_scan term both hit 80s 80s_*, *k_scan term both hit 80s 40s_*. These rates relate to the dissociation of scanning ribosomes upon collisions with a 5’ elongating ribosome. Both hit refers to collisions with ribosomes on both sides. In the collision-mediated 40S dissociation model, the following rates are equal and non-zero: *k_scan term 3 hit 40s_*, *k_scan term 3 hit 80s_*, *k_scan term both hit 40s 40s_*, *k_scan term both hit 40s 80s_*, *k_scan term both hit 80s 40s_*, *k_scan term both hit 80s 80s_*. These rates relate to the dissociation of scanning ribosomes upon collisions with a 3’ scanning or elongating ribosome. The *in vivo* abortive termination rates of scanning ribosomes are not known. Small ribosomal subunits that make it to the 3’ end of the mRNA through leaky scanning of all (u)ORFs always dissociate.

### Model calibration to reporter measurements

We derive the *k_cap bind_* rates by spline interpolation of computationally modeled protein output fit to experimental data (Fig. 2C). We minimized the root mean square error between modeled protein output across variations in these parameters and the experimental data.

### Human uORF search

We import uORF lists from several databases^6,76,77^. The SmProt database77 includes 3162 uORFs from ribosome profiling data, which we filter down first to 1080 uORFs after filtering for aligned matches, available Kozak context, near-cognate start codons, and non-duplicates. Two of these uORFs end in diproline motifs, including *C1orf43*. Another database is a set of high confidence ORFs derived from ribosome profiling of human-induced pluripotent stem cells (iPSCs) or foreskin fibroblast cells (HFFs) and was downloaded from https://www.ncbi.nlm.nih.gov/pmc/articles/PMC4720255/bin/NIHMS741295-supplement-3.csv^6^. This database includes 1517 high confidence (ORF-RATER score > 0.8) uORFs from either iPSCs or HFFs, which we filter down to 3 that end in diproline motifs, including *ABCB9, C1orf43*, and *TOR1AIP1*. The third database derives from HEK293T, HeLa, and K562 cells using ribosome profiling and was downloaded from https://static-content.springer.com/esm/art%3A10.1038%2Fs41589-019-0425-0/MediaObjects/41589_2019_425_MOESM3_ESM.xlsx^76^. This database includes 3577 uORFs which we filter down to 3 that end in diproline motifs and that are less than 60 residues in length for ease of cloning, including *ABCB9,C15orf59*, and *PPP1R37*.

---

**Figure S1.**
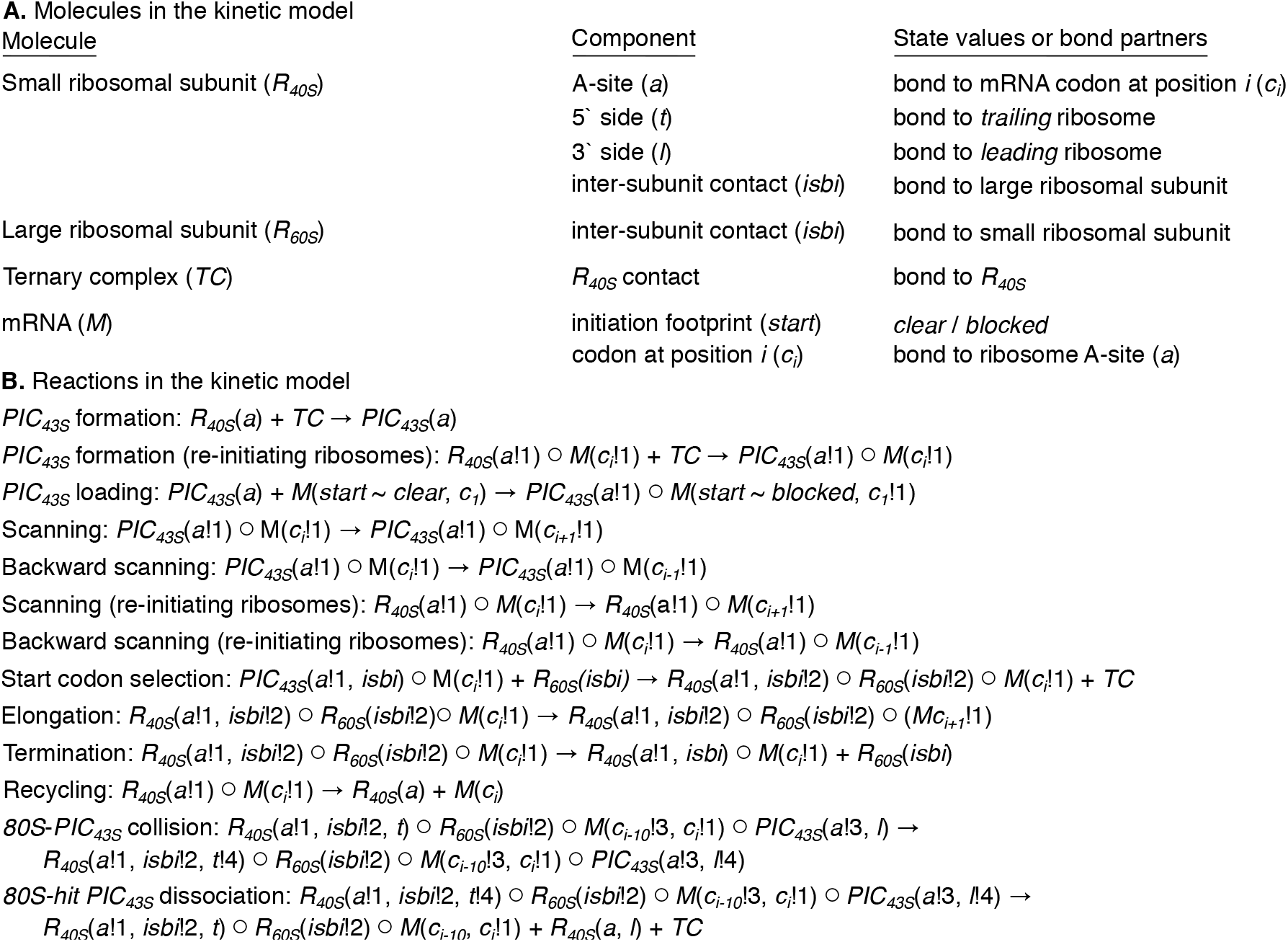
Modeling workflow. **(A)** Molecules in the kinetic model. Molecules have components each of which has state values or bond partners. For example, the mRNA (*M*) initiation footprint (*c_1_* to *c_n_* where n is equal to the ribosome footprint size in nt) can either be *clear* of ribosomes, and therefore free to allow a *PIC_43S_* loading reaction, or *blocked* by a ribosome and unable to allow this reaction. **(B)** Reactions in the kinetic model. Exclamation points and the following numbers indicate component interactions following BioNetGen convention^53^. For example in the *PIC_43S_* loading reaction, the A-site (*a*) of the *PIC_43S_* binds to the mRNA (*M*) at codon 1 (*c_1_*, the 5’ cap). These interactions create the molecule bindings indicated by open circles. Plus signs between molecules indicate that they are not bound together. Re-initiation necessitates several additional reactions. *PIC_43S_* formation (*R_40S_* binding *TC*) can occur if the *R_40S_* is bound to the mRNA; this *TC* re-binding is required for start codon selection competence. *R_40S_* molecules can forward and backward scan. Some reactions in the kinetic model, namely different types of collision and dissociation reactions, are not depicted here.

**Figure S2.**
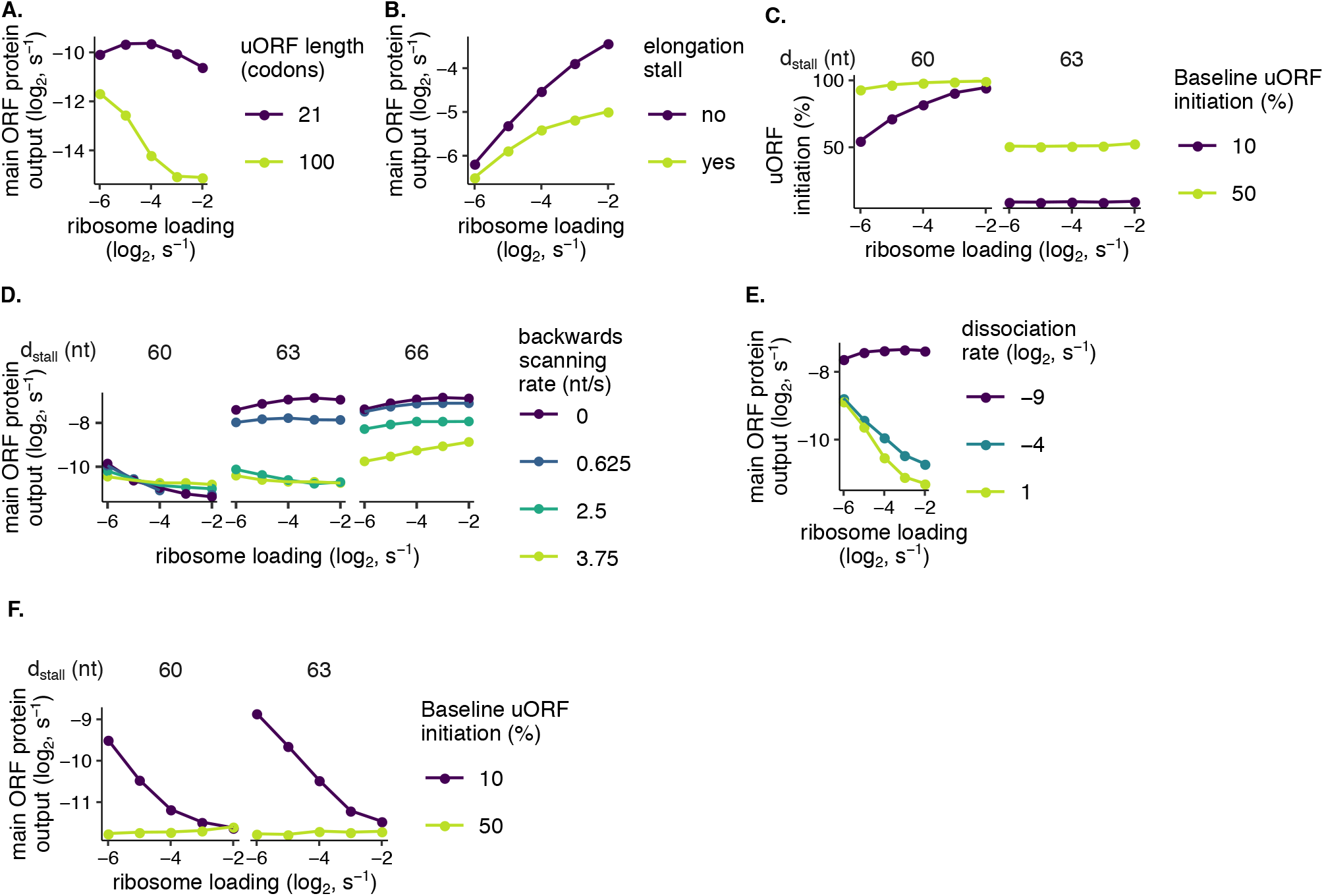
**(A)** Buffering in the 80S-hit dissociation model is affected by uORF length. Re-initiation is 0.2%. uORF initiation is 80%. **(B)** Buffering in the 80S-hit dissociation model is lost with control matched parameters. Buffering in the 80S-hit dissociation model requires strong uORF initiation and rare re-initiation (Fig. 3B, left panel, yellow-green line) and is stronger with longer uORFs (Fig. S2A, yellow-green line). However, we estimate re-initiation to be frequent (Table 1) following calibration of our modeling (Fig. 2D) to reporter measurements on wild-type or mutant uORF2 (Fig. 2C). uORF initiation is 2%. When the elongating ribosome stall is present, *d_stall_* is 63 nt to prevent reduction to the queuing-mediated enhanced repression model. **(C)** Queuing-mediated enhanced uORF initiation is sensitive to *d_stall_*. As the rate of ribosome loading increases, the average queue size increases and allows enhanced uORF initiation only when *d_stall_* equals an integer multiple of the ribosome footprint (30 nt). **(D)** Backward scanning only relaxes the depending of buffering on *d_stall_* in the queuing-mediated enhanced repression model when *d_stall_* is close to an integer multiple of the ribosome footprint (30 nt). The forward scanning rate is 5 nt/s. For *d_stall_* values of 60, 63, 66 nt, the uORF length is 21, 22, 23 codons, respectively. **(E)** Buffering in the collision-mediated 40S dissociation model occurs even with a rather low dissociation rate. Here, *d_stall_* is 63 nt. **(F)** Buffering in the collision-mediated enhanced repression model (Fig. 1D) is insensitive to *d_stall_*. All rates and labels are identical to the main modeling figure unless otherwise specified. Error bars of simulated data are smaller than data points.

**Figure S3.**
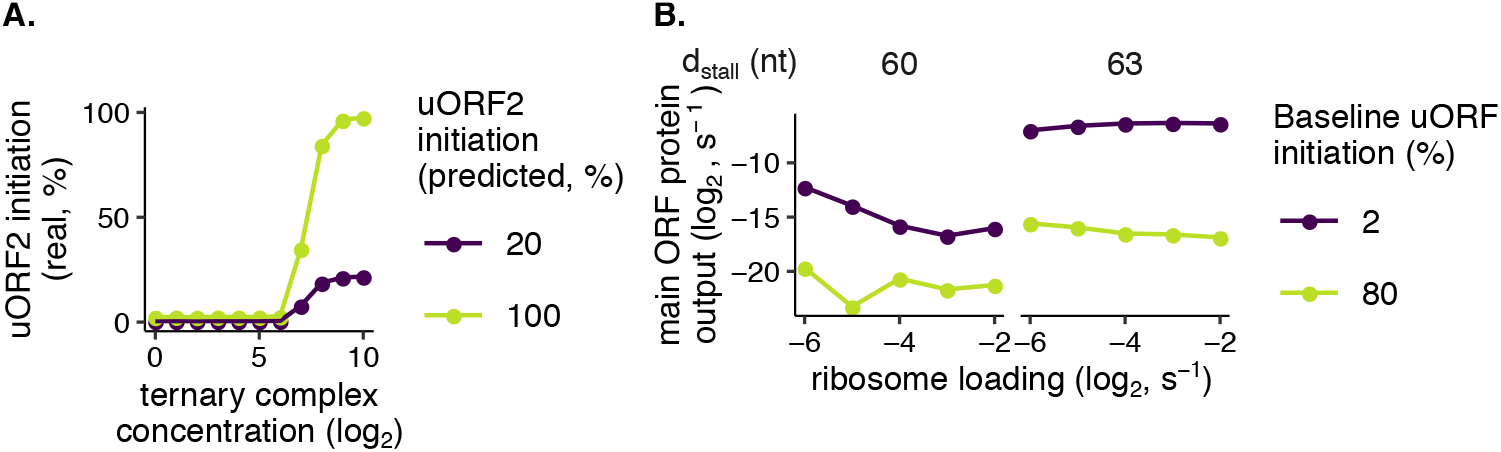
**(A)** Initiation at the second downstream uORF is dependent on high ternary complex concentration. Initiation at the first uORF is 100%. Continued scanning fractions at both uORFs are 100%. Following termination at the first uORF, initiation at the second downstream uORF depends on if a new ternary complex has been acquired since termination at the first uORF. Only when ternary complex concentration is high does this real uORF2 initiation fraction approach the predicted fraction. **(B)** With an elongating ribosome stall, the 80S-hit dissociation model acquires *d*_stall_-dependent buffering similar to that in the queuing-mediated enhanced repression model (Fig. 3C). Re-initiation is 0.2%. All rates and labels are identical to the main modeling figure unless otherwise specified. Error bars of simulated data are smaller than data points.

**Table S1.**
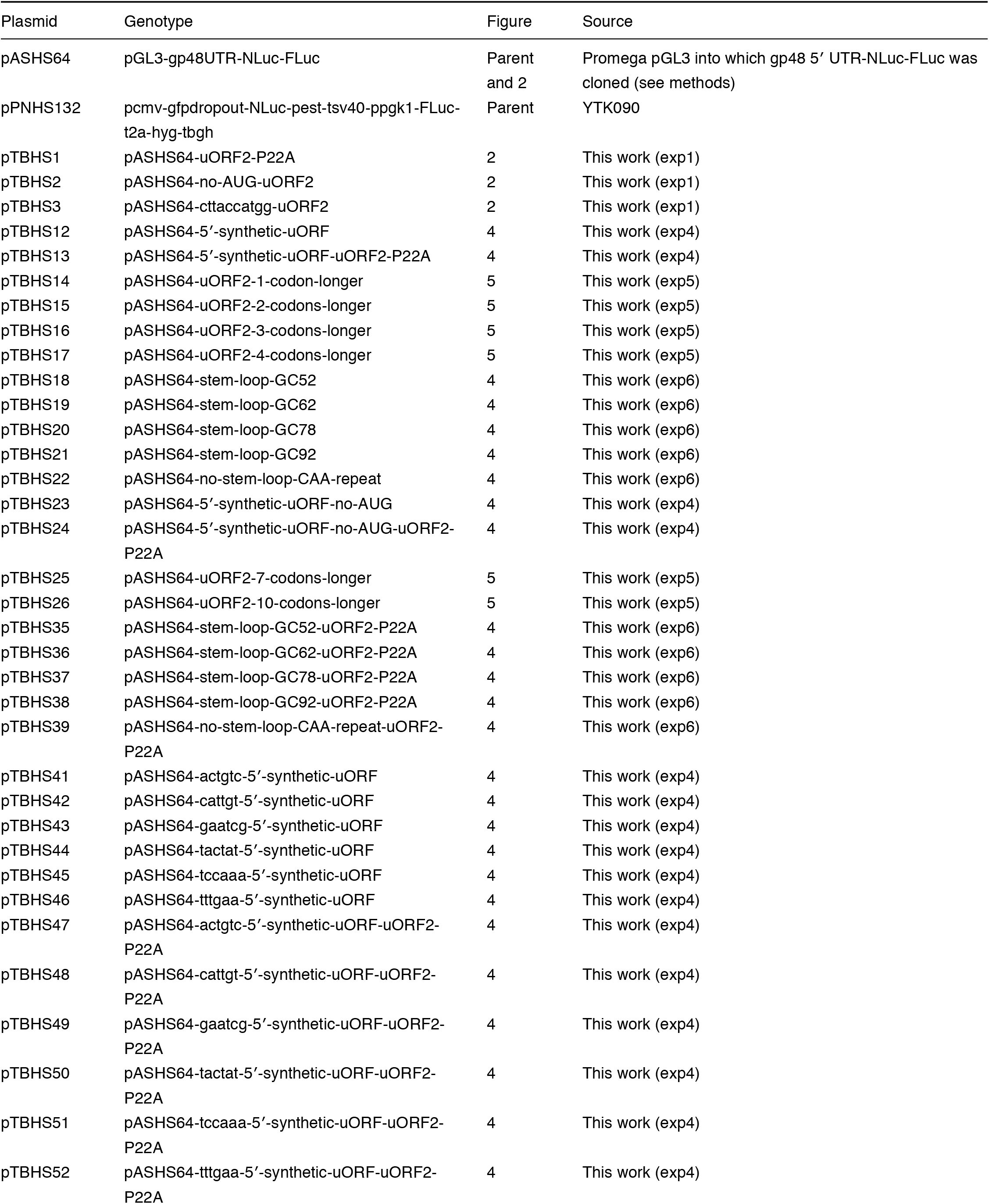
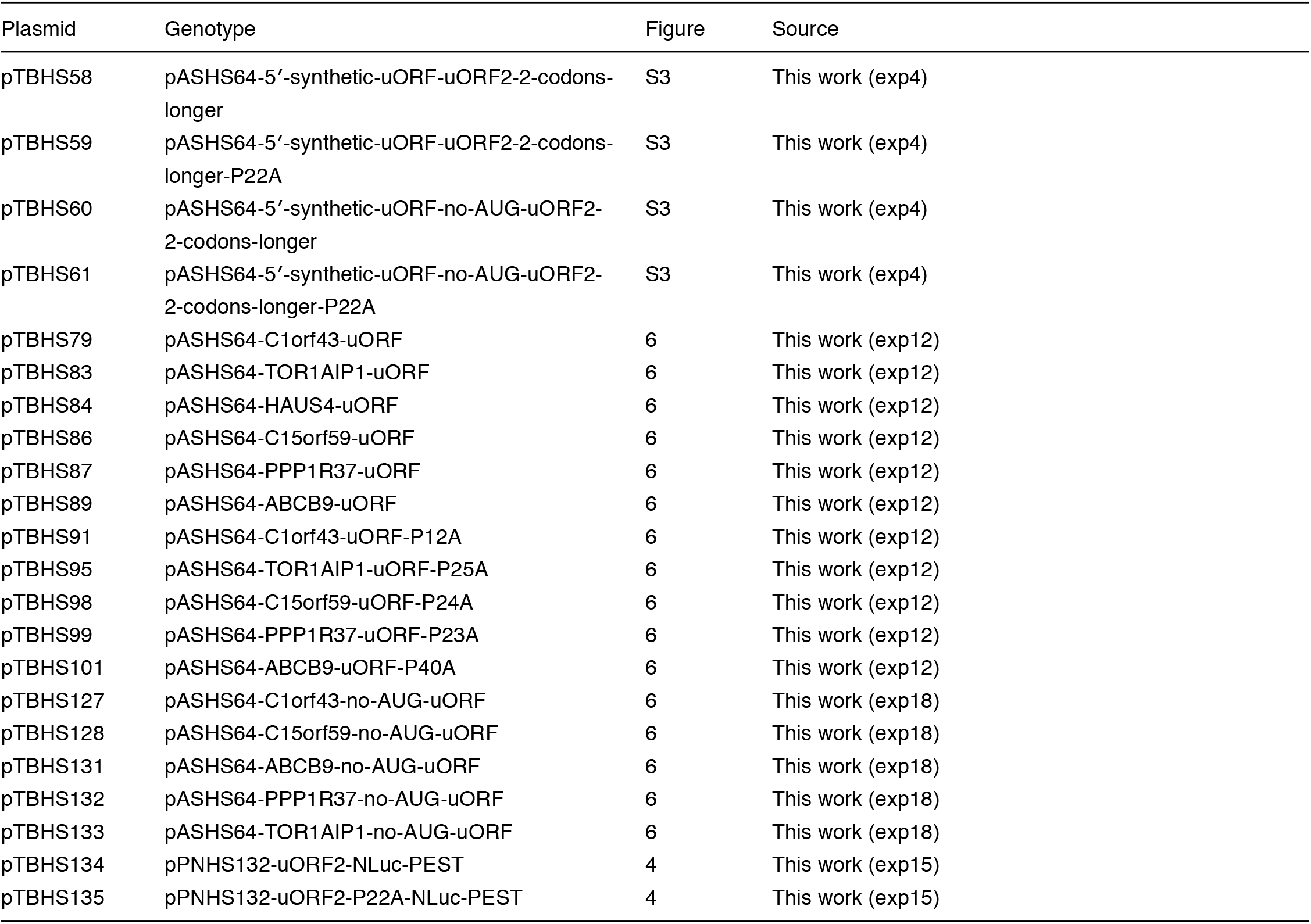
List of plasmids used for this study.

